# An intergenic non-coding RNA promoter required for histone modifications in the human β-globin chromatin domain

**DOI:** 10.1101/639807

**Authors:** Emmanuel Debrand, Lyubomira Chakalova, Joanne Miles, Yan-Feng Dai, Beatriz Goyenechea, Sandra Dye, Cameron S. Osborne, Alice Horton, Susanna Harju-Baker, Ryan C. Pink, Daniel Caley, David R. F. Carter, Kenneth R. Peterson, Peter Fraser

## Abstract

Transcriptome analyses show a surprisingly large proportion of the mammalian genome is transcribed; much more than can be accounted for by genes and introns alone. Most of this transcription is non-coding in nature and arises from intergenic regions, often overlapping known protein-coding genes in sense or antisense orientation. The functional relevance of this widespread transcription is unknown. Here we characterize a promoter responsible for initiation of an intergenic transcript located approximately 3.3 kb and 10.7 kb upstream of the adult-specific human β-globin genes. Mutational analyses in β-YAC transgenic mice show that alteration of intergenic promoter activity results in ablation of H3K4 di- and tri-methylation and H3 hyperacetylation extending over a 30 kb region immediately downstream of the initiation site, containing the adult δ- and β-globin genes. This results in dramatically decreased expression of the adult genes through position effect variegation in which the vast majority of definitive erythroid cells harbor inactive adult globin genes. In contrast, expression of the neighboring ε- and γ-globin genes is completely normal in embryonic erythroid cells, indicating a developmentally specific variegation of the adult domain. Our results demonstrate a role for intergenic non-coding RNA transcription in the propagation of histone modifications over chromatin domains and epigenetic control of β-like globin gene transcription during development.

## Introduction

A staggering proportion of the mammalian genome is transcribed. More than half of the transcribed genomic regions in mammalian cells are intergenic regions [1–3] producing vast amounts of non-coding RNAs [4]. Such intergenic transcription patterns can be complex, with multiple overlapping sense, and antisense transcripts detected for a single locus [5]. During the process of mammalian development, proper temporal and spatial expression of genetic information must be achieved. The multigene β-globin locus has been intensively studied as a model for these events. In humans, it spans over 70 kb on chromosome 11 and its five genes (*HBE*, *HBG1*, *HBG2*, *HBD* and *HBB*) are arranged in the order of their developmental expression in erythroid cells. The locus includes a locus control region (LCR) [6], located 6-22 kb upstream of the embryonic *HBE* gene, which acts as a long-range regulatory element that physically interacts with the active globin genes during transcription [7, 8].

Our previous studies [9] have shown that the locus is divided into at least three differentially activated and developmentally regulated chromatin subdomains of 20-30 kb each (the LCR, εγ, and δβ domains). Each domain displays differential general sensitivity to DNAse I and extensive developmentally regulated H3K4 di and tri-methylation (H3K4me2 and H3K4me3) and H3 hyperacetylation [10]. Each of the domains are also delineated by non-coding intergenic transcription [9], similar to the *Hox* clusters [11, 12], correlating with their differential nuclease sensitivity and active histone modifications. Intergenic transcripts initiating from multiple discrete sites can be found in or near the LCR [13–16] at all stages of development in erythroid cells [9, 17], consistent with a developmentally stable open chromatin conformation. Intergenic transcripts in the gene domains appear during development in coordination with active expression of the globin genes within the embryonic or adult domains. In the case of the adult-specific δβ domain, 5’ RACE has shown that intergenic transcripts initiate near the 5’ boundary of the DNAse I sensitive domain [9]. The analysis of transgenic mice carrying a human β-globin locus yeast artificial chromosome (β-YAC) with a 2.5kb deletion [18] encompassing the intergenic transcript initiation site suggested that the region is necessary for the normal DNase I sensitive chromatin domain surrounding the adult genes and high-level, non-variegated expression of *HBD* and *HBB*. Interestingly, a long-range chromatin interaction between the LCR and this region was recently revealed by 5C analysis of the *HBB* locus, and analysis of the various naturally occurring deletion of this region, results in defective HBG silencing, supporting the suggestion that the intergenic promoter has regulatory potential. In patients carrying the Corfu deletion, *HBG* transcription persists in adult erythroid cells where it effectively competes with the adult *HBB* gene for the activating effects of the LCR [19]. Analysis of intergenic transcripts in Corfu erythroid cells revealed persistent activation of an intergenic promoter upstream of the *HBG* genes (Goyenechea, Chakalova and Fraser, unpublished data) in a region implicated in HbF control. This tight correlation between non-coding transcription domains and domains of increased chromatin accessibility suggested that transcription through chromatin might propagate remodelling or histone modifying activities over large regions. This view is supported by findings of histone acetyltransferases (HATs) and methyltransferases associating with the elongating form of RNA polymerase II and the increasing number of studies linking transcription with simultaneous histone modification. However in other instances the ncRNAs themselves have been implicated in recruiting chromatin modifying activities to chromatin to affect gene expression [12, 20-23]. We previously showed that intergenic transcription in the globin locus is cell cycle-dependent, occurring in S and G1 phases [9, 10], suggesting that it may be involved in the ‘re-establishment’ and/or spreading of an active chromatin state after replication and mitosis.

We hypothesized that regulation of chromatin structural domains plays a key role in control of developmental globin gene expression and that intergenic transcription is required for domain remodeling and subsequent gene activation. To discriminate between cause and effect, as well as effects of other potential regulatory sequences in the larger 2.5kb deletion that was previously analyzed, we generated transgenic mice carrying a 213 kb β-YAC transgene with a 300 bp sequence, corresponding to the putative δβ promoter and transcriptional start site of the adult intergenic transcript, flanked by *loxP* sites. We show that alteration of intergenic promoter activity results in an adult stage-specific reduction in intergenic transcription and a domain-wide, markedly diminished histone modification profile compared to the normal active locus in transgenic mice, leading to dramatically reduced expression of the adult β-like globin genes.

## Results

### Promoter activity in the region upstream of the adult β-globin domain

We previously showed through primer extension and 5’RACE that intergenic transcripts from the 5’ region of the adult-specific β-globin domain appeared to initiate from a discrete site approximately 3.3 kb upstream of the *HBD* gene in transgenic mice [9]. To determine whether sequences in this region have genuine promoter activity, and if so, to perform a preliminary characterization of the promoter elements, we cloned various sub-fragments from –685 to +40 relative to the 5’ end of the transcript in front of a promoter-less EGFP gene (Figure 1A). The resultant plasmids were linearized and stably transfected into the human K562 cell line, which expresses both erythroid and myeloid markers. Populations of transfected cells were assessed for EGFP expression by flow cytometry. The results show that the region indeed has promoter activity (Figure 1B). Expression of EGFP was low but occurred in a significantly higher proportion of transfected cells compared to the promoter-less construct itself or non-transfected cells. The data indicate that most of the promoter activity is contained within the 150-200 bp region immediately upstream of the start site of the intergenic transcript. Truncation down to –50 (Xba) completely abolished promoter activity. The promoter appears to lack a canonical TATAA-like sequence within 30 bp of the start site, although there is an overlapping GATA1 TATAA sequence at approximately –190 bp. However deletion of that sequence from the promoter results in only a slight decrease in EGFP expression (compare Apo to Bst11). In addition, the region from –150 to 0 bp contains putative transcription factor binding sites indicative of possible promoter function. We conclude that the region has promoter activity and that a minimal promoter containing the first few hundred base pairs upstream of the start site retains promoter activity in stably transfected cells.

**Figure 1.**
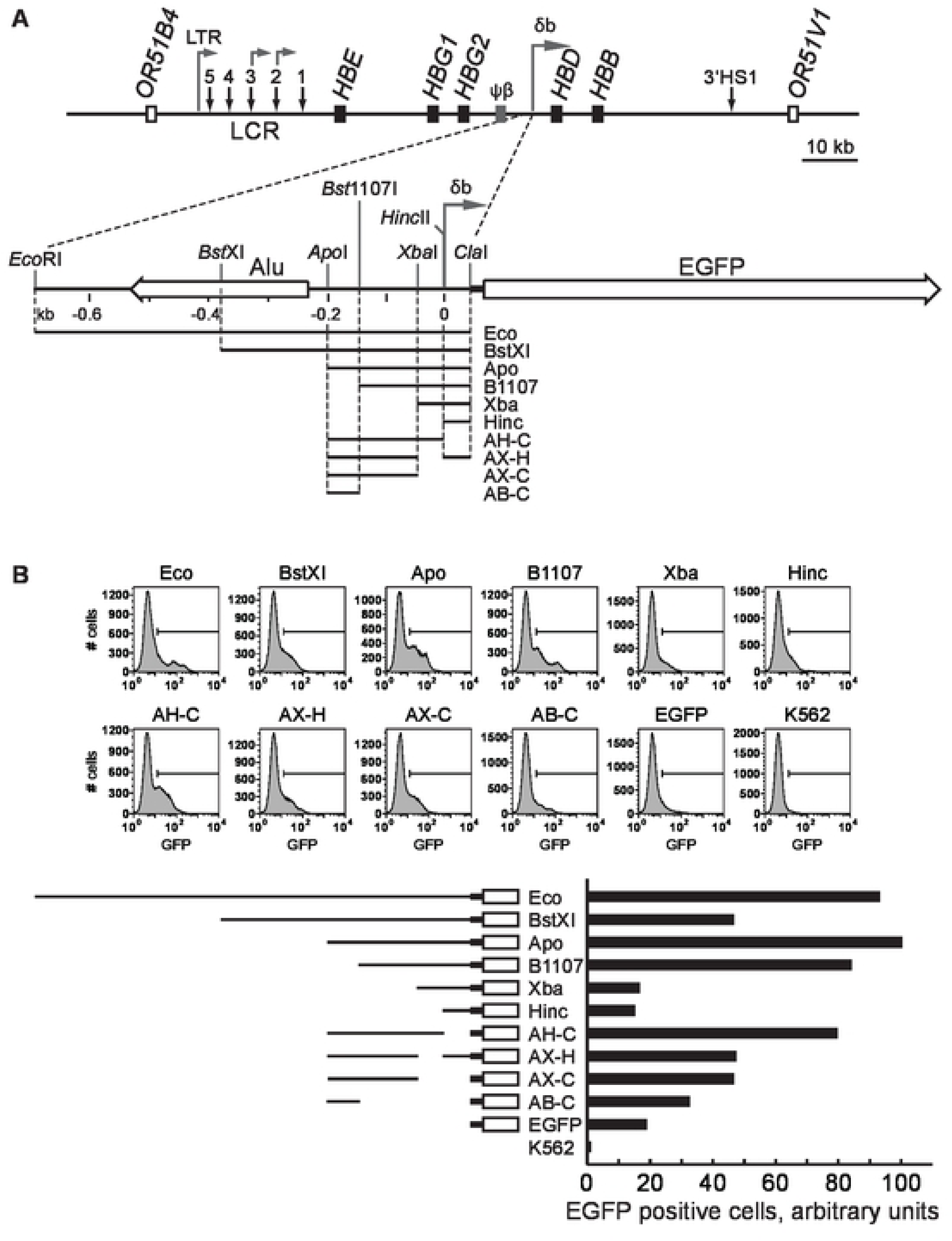
Schematic map of the human β-globin locus and characterization of the δβ intergenic promoter. (**A**) Schematic diagram of the human β-globin locus and the δβ intergenic promoter region. Black boxes represent globin genes; grey box corresponds to the β-like pseudogene, open boxes – to olfactory receptor genes flanking the β-globin locus. Vertical arrows indicate DNase hypersensitive sites, including sites 1 to 5 of the locus control region (LCR); horizontal arrows denote intergenic transcription start sites. The detailed map of the δβ promoter region shows the positions of the intergenic transcription start site and upstream Alu element (left horizontal open arrow). Restriction fragments from the δβ promoter region were cloned upstream of a promoterless EGFP gene as indicated and described in Materials and Methods. Relevant restriction sites are indicated. (**B**) EGFP reporter assays. Constructs were stably transfected into K562 cells and the percentage of EGFP-expressing cells was determined by flow cytometry. The Apo construct, which showed the highest percentage of EGFP-positive cells, was assigned an arbitrary value of 100.

### Analysis of transcription upstream of the adult β-globin domain

To investigate the pattern of intergenic transcription throughout the globin locus in human cells we utilized an *ex vivo* model of erythroid development. Nucleated cells were isolated from human peripheral blood and induced to differentiate in a two-phase culture system. During the second phase of growth the induction of HBB expression was measured and changes were observed in cell morphology characteristic of erythroid differentiation (not shown). We and others have previously shown that this is a good model for studying the activation of the adult β-globin domain [19, 24]. Total RNA was extracted at different time points during this erythroid progression, reverse-transcribed, fluorescently labelled and then hybridized to a custom high-density tiling array. At an early time-point (day 4) we saw no significant peak of transcription over the putative δβ-promoter (Figure 1A), consistent with the relatively lower levels of *HBB* expression and lack of hemoglobinization at this stage of differentiation. Over time we saw an increase in signal over the putative intergenic promoter which peaked at around day 7 of erythroid differentiation (Figure 2A).

**Figure 2.**
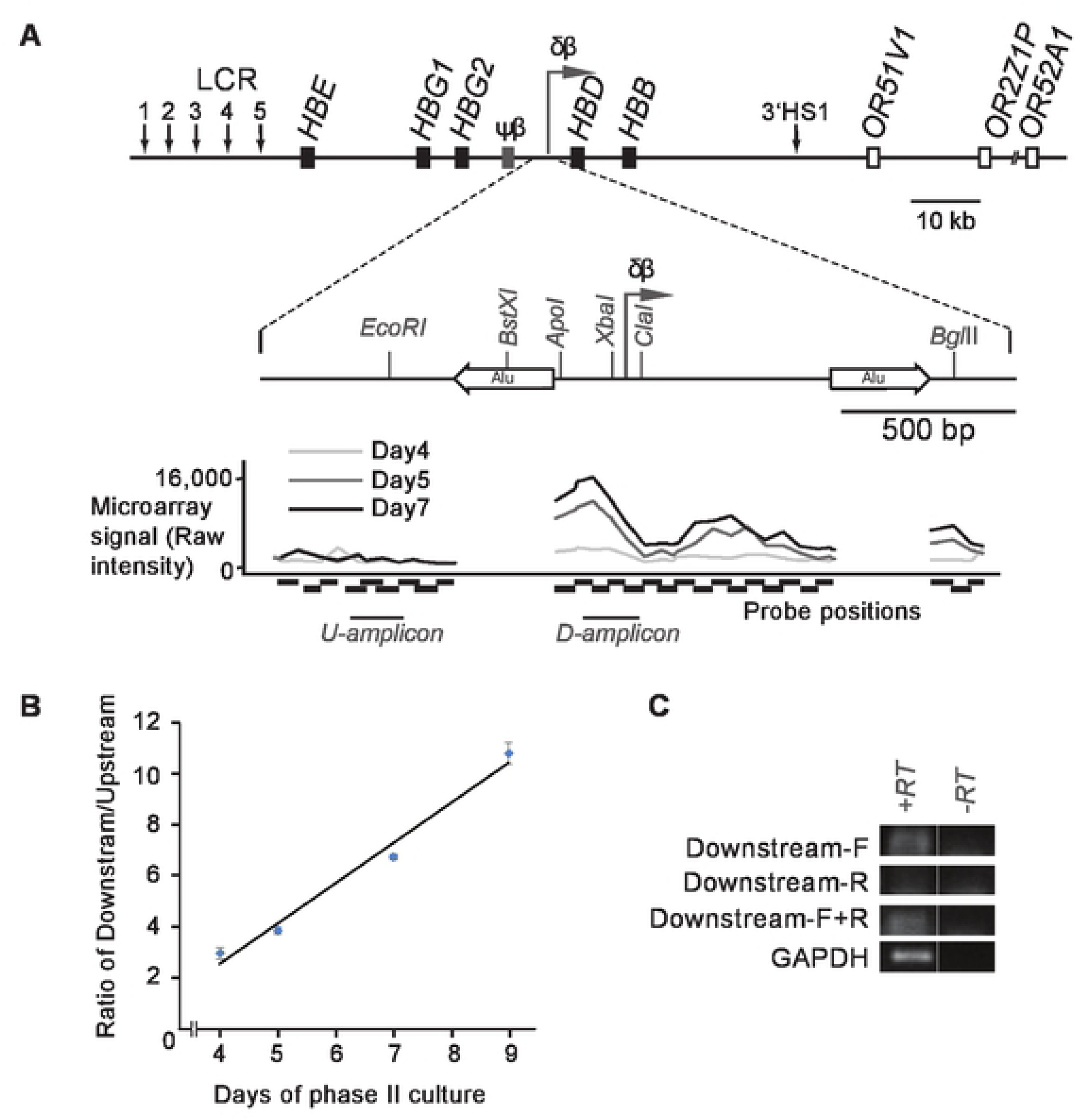
Analysis of transcription in the δβ intergenic promoter region in human erythroid cells. (**A**) Schematic diagram of the human β-globin locus and the δβ intergenic promoter region. Black boxes represent globin genes; grey box corresponds to the β-like pseudogene, open boxes correspond to olfactory receptor genes flanking the β-globin locus. Vertical arrows indicate DNase hypersensitive sites, including sites 1 to 5 of the locus control region (LCR); horizontal arrows denote putative intergenic transcription start sites. The detailed map of the δβ promoter region shows the positions of the intergenic transcription start site and upstream Alu element (left horizontal open arrow). Restriction enzyme sites are indicated by vertical lines. The graph below the δβ intergenic promoter schematic indicates microarray signal intensity for various probes (indicated under the graph with horizontal black boxes) following labelling and hybridisation of RNA extracted from primary erythroid cells at different phases (day 4 in light grey, day 5 in dark grey and day 7 in black) of phase II culture. The results shown are representative of five biological replicates (**B**) The RNA samples used for microarray analysis in A were converted into cDNA using a WTA kit and amplified by PCR. The PCR amplicons are indicated beneath the microarray signal graph as U (upstream) amplicon and D (downstream) amplicon. The amounts of each amplicon were calculated by comparing signals to a standard curve generated with genomic DNA and the ratio of downstream over upstream was calculated. The ratio increases through phase II indicating a relative increase in transcripts downstream of the putative intergenic promoter. Results are representative of two biological replicates. Pearson correlation analysis indicates a statistically significant correlation (p<0.01). (**C**) RNA from day 7 of erythroid differentiation was converted into cDNA using either gene-specific primer (the downstream-amplicon forward primer [Downstream-F], the downstream-amplicon reverse primer [Downstream-R] or a combination of the two primers [Downstream-F+R] as indicated to the left of each panel). PCR was then carried out using both Downstream-F and Downstream-R primers (to amplify the downstream amplicon). GAPDH primers were used as a control to confirm RNA integrity. PCR products were run on a 2% gel, stained with SYBR Gold, and visualised on a Typhoon Imager. The strand-specific results show that the transcripts detected downstream of the δβ intergenic promoter are sense transcripts (travelling towards HBD).

This is consistent with an increase in *HBB* expression at these later stages (not shown). The lack of a peak in the day 4 sample suggests that the peak observed at later time-points is not caused by locally increased non-specific background binding to the probes in this region, but is due to an increase in transcripts in the area. However, to confirm the latter we performed quantitative RT-PCR using primers upstream and downstream of the putative signal. The results show an increase in the ratio of downstream to upstream signal as erythroid differentiation proceeds (Figure 2B), which is consistent with increased transcription from a putative promoter in this region. To determine the directionality of transcription we performed a strand-specific RT-PCR using either Downstream-F (the forward primer, which detects a sense transcript) or Downstream-R (the reverse primer, which detects an antisense transcript) in the cDNA-generation step. Only the Downstream-F primer was able to generate a cDNA that could be amplified in PCR (Figure 2C), suggesting that the transcript downstream of the putative δβ-promoter is fired in the direction towards the adult β-globin sub-domain. Taken together these data suggest that the δβ-promoter is capable of firing transcripts in erythroid cells from humans as well as transgenic mice and reinforces the validity of using the latter as a model to study the role of the δβ-promoter in β-globin regulation.

### Generation of YAC transgenic mice with deletion of the δβ intergenic promoter

To investigate the contribution of the δβ intergenic promoter to the regulation of gene activity and chromatin structure in the globin locus, we designed a 213 kb YAC transgene construct containing the entire human β-globin locus with *loxP* sites flanking the transcriptional initiation site, from –266 to + 40 (Figure 3A). The unmodified YAC has been used extensively in transgenic mice [25–30] and shown to be sufficient for normal, copy number-dependent, position-independent expression of the human beta-globin genes throughout development. The modified YAC was used to generate three transgenic mouse lines using standard microinjection procedures. The structural integrity of the transgene was analyzed by *SfiI* digestion of genomic DNA followed by pulsed field gel electrophoresis and Southern blot using multiple probes across the locus. As can be seen in Figure 3B each of the lines harbors a characteristic large *Sfi*I fragment that hybridizes to all probes from the 5’ end to the 3’ end of the *Sfi*I fragment. This indicates that each line contains at least one intact copy of the *HBB* locus. PCR cloning and sequence analysis showed that lines FX101 and FX115 contained the two flanking *loxP* (flox) sites whereas line FX14 contained only a single *loxP* site located downstream of the δβ promoter. Of the two ‘floxed’ lines we chose FX115 for further study, based on the fact that it contained a single major band containing regions complimentary to all the tested probes verifying the structural integrity of the transgene. This line was bred with a Cre recombinase expressing transgenic line, Deleter Cre [31] to successfully generate a deletion line with a single, intact 115 kb *Sfi*I fragment (Figure 3C). Southern blot (Figure 3D) and PCR analysis (not shown) confirmed that the deletion line Δ115, retains a single copy of the human β-globin locus with the 306 bp floxed δβ promoter region removed.

**Figure 3.**
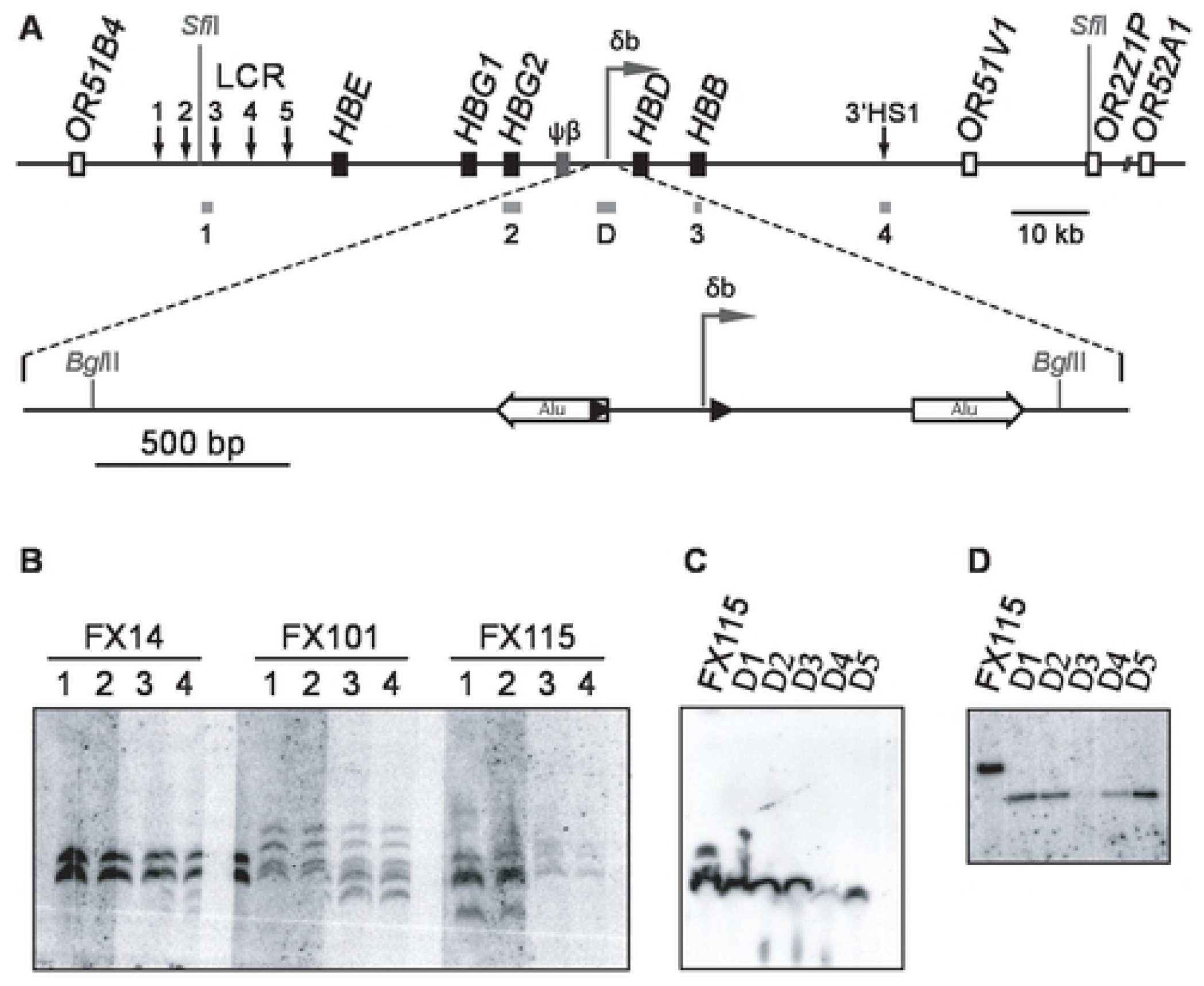
Generation of YAC transgenic mice carrying a floxed δβ intergenic promoter and production of the deletion lines. (**A**) Schematic diagram of the human β-globin locus YAC used to generate transgenic lines FX14, FX101, and FX115. Black boxes represent globin genes; the grey box corresponds to the β-like pseudogene; open boxes indicate olfactory receptor genes. Vertical arrows denote DNase hypersensitive sites. Horizontal arrow refers to the δβ intergenic transcription start site. The δβ promoter region is shown at a larger scale below the main map. Black triangles denote *loxP* sites inserted upstream and downstream of the δβ promoter. White horizontal arrows indicate Alu repeat elements flanking the δβ promoter. *Sfi*I and selected *Bgl*II restriction sites are indicated, and probes are shown as grey boxes below the lines (probes 1 to 4 and probe D described in Materials and Methods). (**B**) YAC transgenic lines carrying the human β-globin locus, before Cre-mediated deletion of the intergenic promoter. Genomic DNA from three transgenic lines, FX14, FX101, and FX115, was digested with *Sfi*I, subjected to pulsed-field electrophoresis and Southern transfer. Several identical DNA samples from each line were run on the same gel, blots were cut into strips and each strip was hybridized with a different probe as indicated. Probes throughout the β-globin locus identify identical *SfiI* fragments. (**C**) Transgenic mice with deleted δβ promoter. Genomic DNA from line FX115 and 5 animals carrying the deletion, Δ1 to Δ5, was digested with *Sfi*I, subjected to pulsed-field electrophoresis, transferred and hybridized with probe D. A single band corresponding to the β-globin *Sfi*I fragment is detected in Δ1 to Δ5. (**D**) Efficient deletion of the δβ intergenic promoter. Southern blot analysis of *Bgl*II-digested genomic DNA from parent line FX115 and Δ1 to Δ5 hybridized to probe D. Note size reduction of *Bgl*II band after deletion of the δβ-promoter.

### Embryonic globin gene expression is unaffected by the δβ promoter deletion

To ascertain the effect of the intergenic promoter deletion on globin gene expression we used RNA FISH to study the transcriptional regulation of the globin genes throughout development. Two wild type YAC transgenic lines with equivalent copy numbers were used for comparison. Line FX14 contains 2 copies of the same human beta-globin YAC as the floxed and deleted lines but has only a single *loxP* site in the position downstream of the δβ promoter. Line 264W contains a single intact copy of a shorter human beta-globin YAC (150 kb) which contains the entire globin locus. This line has previously been shown to express the *HBB* genes at full levels, equivalent to the endogenous mouse *Hbb* genes [32] and has normal patterns of intergenic transcription and histone modifications [10]. Transgenic mice containing the entire human globin locus and intact LCR normally express *HBE* and *HBG* at high levels in E10.5 embryonic red cells. RNA FISH analysis shows that embryonic expression of *HBE* and *HBG* is normal for lines FX115 and Δ115 (Figure 4A) with approximately 90% of the embryonic cells being positive for *HBE* and/or *HBG* transcription signals (Figure 4B). This degree of transcriptional activity is equivalent to the two wild type lines 264W and FX14, and previously published full-locus transgenic lines [33, 34], as well as the endogenous mouse *Hbb* locus in embryonic blood (>90%) [35]. Line FX 101 showed a similar high-level of transcription signals (>90%) in embryonic blood cells (data not shown). These results demonstrate that transcription of the human *HBE* and *HBG* genes is normal and non-variegated in these lines and is not affected by the intergenic δβ promoter deletion, implying position independent transgene expression provided by a fully functioning LCR.

**Figure 4.**
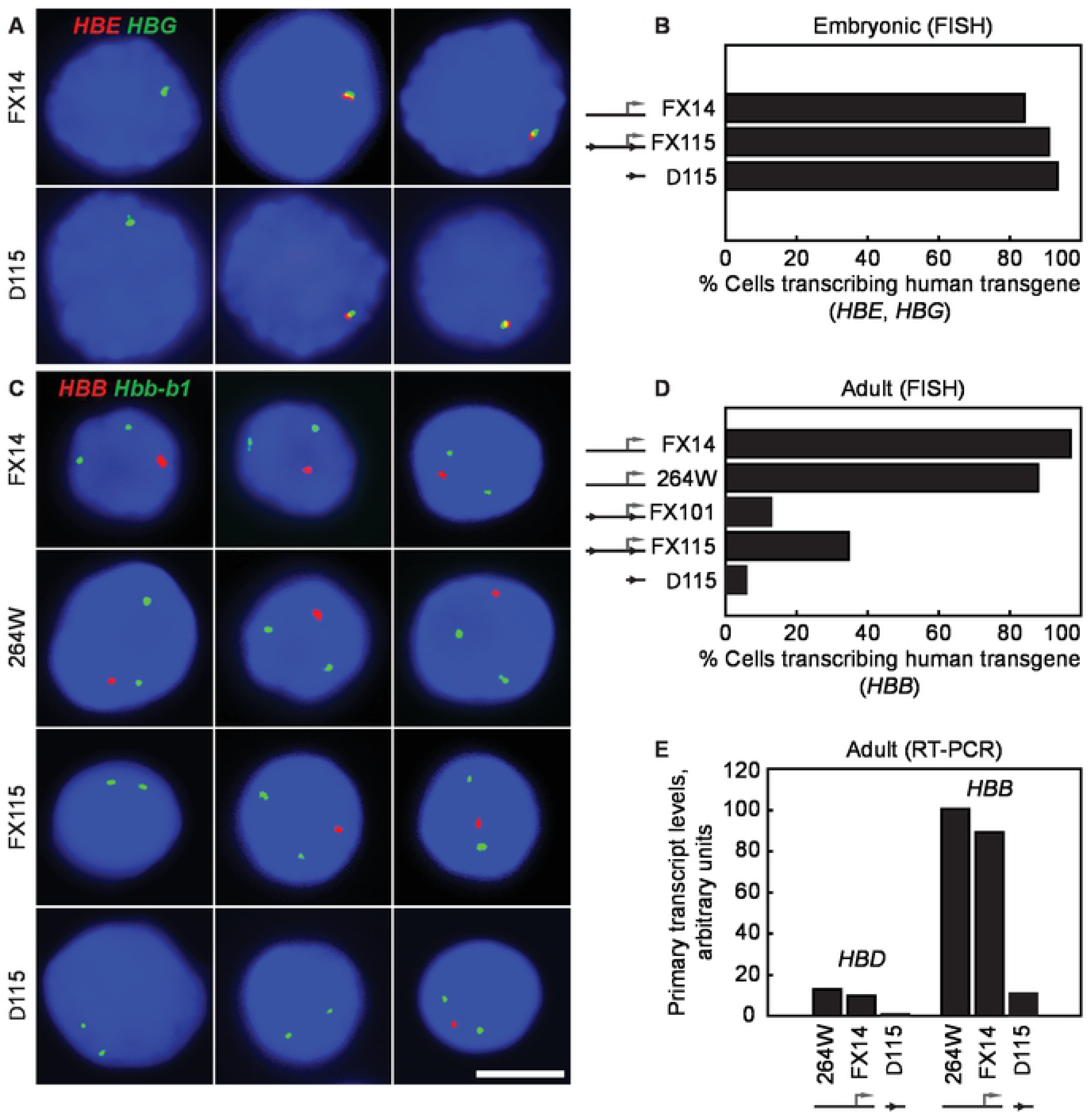
Human β-globin transcription in transgenic mice before and after deletion of the δβ intergenic promoter. (**A**) RNA FISH analysis of human *HBE* and *HBG* transcription in E10.5d blood cells from heterozygous transgenic embryos. *HBE* primary transcripts are in red, *HBG* transcripts in green; for scale, see (C), scale bar, 5 μm. (**B**) Normal transcription of *HBE* and *HBG* genes in E10.5d embryonic blood cells by RNA FISH. Cells positive for human globin gene transcription signals were scored; in each line nearly 100% of the embryonic blood cells have transcription signals for *HBE*, *HBG* or both genes indicating normal high-level expression. (**C**) RNA FISH analysis of *HBB* transcription in adult erythroid cells. *HBB* primary transcripts are in red, mouse *Hbb-b1* transcripts detected in green as an internal control to identify erythroid precursors; scale bar, 5 μm. (**D**) Reduced transcription of *HBB* in adult erythroid cells carrying a δβ promoter deletion. Heterozygous mice from wild-type lines FX14 and 264W, lines FX101 and FX115 with ‘floxed’ δβ promoter, and Δ115 with deleted δβ promoter were analyzed by RNA FISH. Nuclei with a *HBB* signal were scored and expressed as a percentage of *Hbb-b1* positive nuclei. *HBB* transcription is impaired in lines FX101 and FX115, and further reduced in line Δ115. (**E**) Reduced transcription of human *HBD* and *HBB* genes in adult erythroid cells carrying a δβ promoter deletion. Animals from wild-type lines FX14 and 264W, and line Δ115 were analyzed by real-time RT-PCR, as described in Materials and Methods. Primary transcript levels are corrected for transgene copy number; wild-type *HBB* level in 264W is assigned an arbitrary value of 100.

### Intergenic promoter deletion severely impacts adult globin gene transcription

The results in adult erythroid cells were strikingly different from the embryonic globin gene transcription pattern. Unexpectedly, we found that transcription of the adult-specific *HBB* gene was reduced 2-3 fold in the FX101 and FX115 lines in which the *δβ* intergenic promoter is still present with flanking *loxP* sites (Figure 4C and D). Approximately 35% (34.6 %; n=321) of the erythroid cells in FX115 were positive for a human β-globin transcription signal compared to >90% in the wild type lines (see below). Strikingly, we found that deletion of the intergenic promoter (line Δ115) further reduced transcription of *HBB* compared to FX115. Only 5.8% (n=431) of the erythroid cells were positive for an *HBB* transcription signal in Δ115, approximately 6-fold lower than FX115 and 15-20 fold lower than the wild type lines. *HBB* transcription in both wild type lines were similar to the endogenous *Hbb* locus, in which >90% of β-globin loci are positive for transcription signals [35]. The dramatic reduction in adult-specific transcription of *HBB* in the intergenic promoter deletion line Δ115 shows that the 300bp sequence contains an element essential for normal high-level expression of the adult β-globin genes. This was confirmed by RT-PCR analysis, which showed that expression of both adult-type beta-genes; *HBB* and *HBD* are dramatically reduced in the deletion line compared to the wild type and floxed lines (Figure 4E). These results suggest the minimal intergenic promoter is necessary for high-level expression of both *HBB* and *HBD*, suggesting a possible domain-wide effect.

### Flanking l*oxP* sites interfere with intergenic promoter activity

An unexpected finding from the above experiments was the fact that both floxed lines FX101 and FX115 showed reduced transcription of the adult-specific *HBB* gene even prior to deletion of the intergenic promoter (Figure 4D). This raised the question of potential structural problems in our YAC transgene, which may have been introduced, for example, during the process of homologous recombination in yeast. Pulsed-field gel analysis indicated that each of the YAC lines in question contained an intact *Sfi*I fragment that hybridized with several probes across the locus suggesting that the bulk of the β-globin locus was intact (Figure 3B and C). Southern blot analyses using probes around the δβ promoter did not reveal any obvious rearrangements (data not shown and Figure 3D). Similarly, sequence analysis of a 3.3 kb PCR product amplified from genomic DNA from line FX115 showed that the genomic sequence spanning the entire targeted region was the same as wild type except for the insertion of the *loxP* sites, thus ruling out rearrangements in that area during homologous recombination. We then considered the possibility that the inserted *loxP* sites interfered with intergenic promoter activity. To test this we measured intergenic transcript levels downstream of the δβ promoter in FX115 and Δ115 via RT-PCR. We found that both lines had reduced intergenic transcript levels compared to the wild type transgenic strains 264W and FX14 (Figure 5A), which have similar transgene copy numbers. These results suggested that the *loxP* sites may indeed be affecting intergenic promoter activity in FX115.

**Figure 5.**
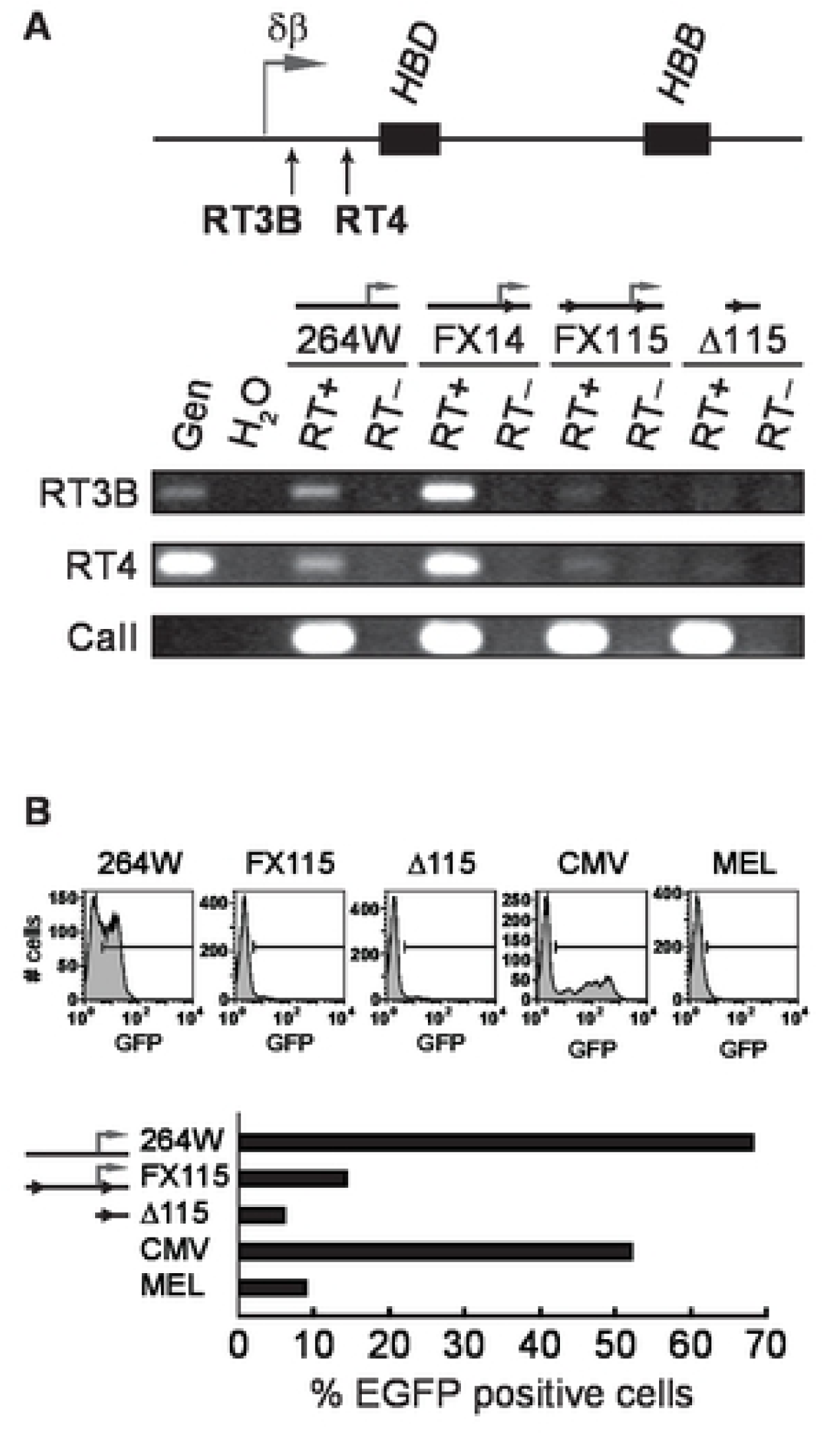
Intergenic transcription in the β-globin locus before and after deletion of the δβ intergenic promoter. (**A**) RT-PCR to detect intergenic transcripts downstream of the δβ intergenic promoter in adult erythroid cells. The positions of PCR amplicons RT3B and RT4 are indicated below the map; black boxes, *HBD and HBB* genes; horizontal arrow, normal position of the δβ intergenic promoter and transcript start site. Intergenic transcription is reduced in lines FX115 and Δ115 with ‘floxed’ and deleted δβ intergenic promoter, respectively. (**B**) Transfection assays to assess intergenic promoter activity. Fragments containing the δβ intergenic promoter or corresponding fragments from the deletion lines were PCR amplified from lines 264W, FX115, and Δ115, and cloned upstream of a promoterless EGFP gene as described in Materials and Methods. Reporter constructs were stably transfected into MEL cells and the number of EGFP-expressing cells in the population was determined by flow cytometry. The percentage of cells expressing EGFP is severely decreased compared to wild-type δβ promoter (assigned an arbitrary value of 100) and CMV promoter when driven by FX115- and Δ115-derived sequences.

To further investigate this possibility we used genomic DNA from FX115 and Δ115 to amplify PCR fragments containing the intergenic promoter regions out to approximately –700 relative to the intergenic transcription start site. These fragments were cloned in front of a promoter-less EGFP reporter plasmid with a flanking neo resistance gene. The plasmid constructs were sequenced to verify that no mutations occurred during the amplification or cloning procedures, and were then stably transfected into mouse erythroleukemia (MEL) cells. EGFP expression was measured in neo-resistant, transfected populations by flow cytometry. The wild type δβ intergenic promoter drove consistent, low-level expression of the EGFP reporter in a very high percentage (66%) of stably transfected cells, compared to the potent CMV promoter/enhancer combination that we used as a positive control (Figure 5B). The CMV promoter population had only 51% EGFP positive cells but the level of expression per cell varied much more widely and was in general much higher than the δβ promoter. As expected, the Δ115 promoter region with the 300 bp deletion showed no EGFP expression being essentially identical to mock transfected cells. Interestingly, the δβ intergenic promoter with flanking *loxP* sites from FX115 had dramatically reduced EGFP expression, only slightly higher than the promoter deletion construct. These results show that the presence of the *loxP* sites partially interferes with intergenic promoter activity by dramatically decreasing the percentage of cells that achieve expression. Deletion of the 300 bp region leads to more or less complete abolition of promoter activity. These results provide a potential explanation for the reduced percentage of cells with *HBB* transcription signals in FX115 and FX101. We conclude that δβ promoter activity and intergenic transcription are essential for normal high-level activation of the adult β-globin genes.

### Absence of intergenic transcription results in variegated transgene expression

In order to determine whether the absence of transcription of *HBB* in most erythroid cells is due to an instability of transcriptional activity, i.e. oscillating between the ON and OFF states in every cell, or the result of variegation of expression in which only a subset of cells stably express the *HBB* transgene, we performed RNA FISH with exonic *HBB* probes which detect both primary transcripts in the nucleus and processed *HBB* mRNAs in the cytoplasm[34]. We observed that only the erythroid cells exhibiting a *HBB* primary transcript signal in the nucleus had spliced *HBB* mRNA transcripts in the cytoplasm and vice-versa (Figure 6, and Figure 4D). This demonstrates that disruption of δβ promoter activity leads to developmental stage-specific variegation of the *HBB* gene in erythroid cells, in which the vast majority of erythroid cells (∼90% in Δ115) never switch on or activate the human beta-globin transgene locus [33]. These findings reinforce the concept that the δβ intergenic promoter element is essential to confer the correct epigenetic state of the adult domain containing the *HBD* and *HBB* genes.

**Figure 6.**
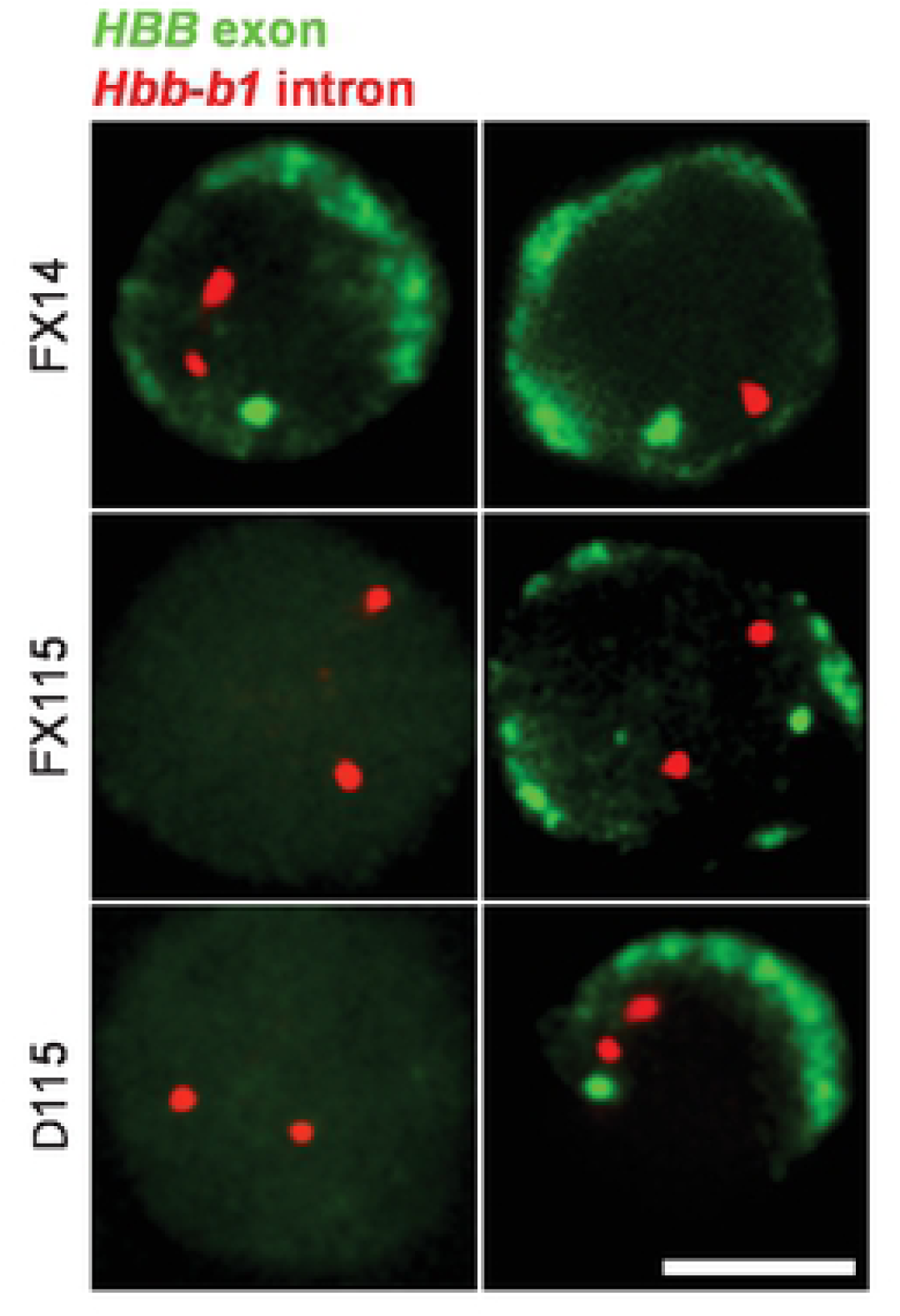
*HBB* expression is variegated in transgenic mice carrying a deletion of the δβ intergenic promoter. RNA FISH analysis of *HBB* transcripts in adult erythroid cells; primary and mature *HBB* transcripts are detected in green by an exon probe; as an internal control, *Hbb-b1* primary transcripts are detected in red to identify erythroid cells; scale bar, 5 μm. Greater than 95% of FX14 erythroid cells have nuclear primary *HBB* signals and cytoplasmic *HBB* mRNA staining (top row). Conversely, the majority of FX115 and Δ115 cells lack primary *HBB* signals and do not contain cytoplasmic *HBB* mRNA (cells on the left). The few FX115 and Δ115 cells that do express *HBB* mRNA in the cytoplasm also have nuclear *HBB* primary transcript signals and vice-versa (cells on the right) showing that *HBB* expression is restricted to a small subset of adult erythroid cells.

### Absence of intergenic transcription results in altered histone modification profile

We previously showed that YAC transgenes with a much larger (2.5 kb) δβ promoter deletion resulted in a 2- to 3-fold reduction in general DNase I sensitivity over the entire adult chromatin domain indicating that the normal, stage-specific chromatin opening of the domain surrounding the *HBD* and *HBB* genes failed in the absence of intergenic transcription [9]. In most cases studied, a DNase I sensitive chromatin conformation correlates with the appearance of “active” epigenetic modifications to histone tails in the region. Using both human primary erythroid cells and erythroid cells from transgenic mice 264WT at different developmental stages, we identified developmentally specific sub-domains of histone H3 hyper-acetylation and H3K4 di- and tri-methylation that correlate with the occurrence of intergenic transcription and increased general DNAse I sensitivity in the human beta-globin locus [10]. Having shown that deletion of the 300 bp core intergenic promoter results in severe variegation of the *HBB* and *HBD* genes, we sought to investigate the effect on histone modifications over this domain. We performed chromatin immunoprecipitations (ChIP) with antibodies directed against histone tail modifications characteristic of active chromatin followed by real time PCR to assess the effect of the deletion on the histone modification profile of the transgene locus in adult erythroid cells.

The ChIP data shows that deletion of the δβ promoter results in a dramatic reduction of H3K4 di- and tri-methylation and H3 hyper-acetylation across the entire adult domain (Figure 7), from approximately 3.5 kb upstream of the *HBD* gene, extending over 30 kb downstream to 3’ HS1 which may correspond to the 3’ boundary of the adult domain [36]. The decreases ranged from 2 to 10 fold compared to the normal *HBB* transgene locus (264W) [10]. Reduction in histone modifications were not restricted to the gene promoters but occurred over intergenic regions throughout the domain. These results show that the spreading of active histone modifications across the entire adult domain is not simply dependent on the presence of intact gene promoters and the LCR, but requires the δβ intergenic promoter.

**Figure 7.**
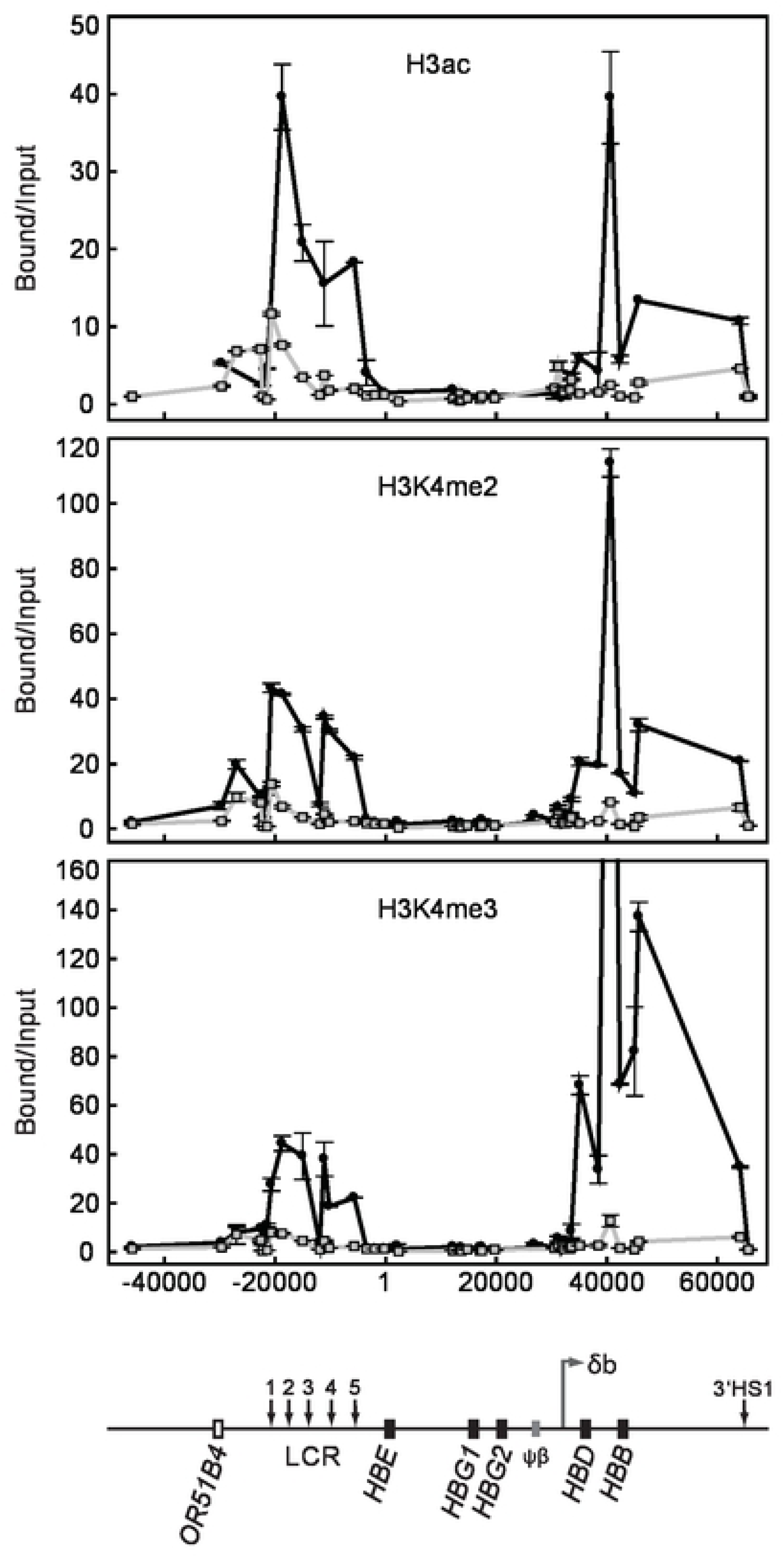
Histone modifications across the human β-globin locus in wt and δβ promoter deleted adult erythroid cells. Chromatin immunoprecipitation assays for histone H3 acetylation (H3ac), di-methylated lysine 4 of histone H3 (H3K4me2), and tri-methylated lysine 4 of histone H3 (H3K4me3); 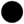, wild-type adult erythroid cells from line 264W [10]; 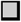, adult erythroid cells from line Δ115. Coordinates in bp are shown along the x axis with *HBE* start site as +1. Bound versus input ratios were calculated and normalized to the most 3’ primer pair, located just downstream of 3’HS1 in the ORG cluster. A map of the locus is shown below the graphs.

A surprising finding from these experiments was that the δβ promoter deletion also resulted in a significant reduction in active histone modifications throughout the LCR domain. The LCR was initially described as a dominant control element capable of chromatin activation at any integration site in the genome. Those early studies used constructs with genes directly linked to the LCR rather than at long range and normal domain context. Our results show that the LCR, though fully active in embryonic cells, is not capable of maintaining an active profile in adult erythroid cells in the absence of an active gene domain within the locus. RNA FISH in Δ115 embryonic and early fetal liver cells (Figure 4 A, B and not shown) shows normal expression of the embryonic globin genes *HBE* and *HBG*, implying full LCR function in primitive and early definitive erythroid cells. Normal, high level expression of these genes was confirmed by RT-PCR analysis (not shown). These results suggest some form of activation cooperativity or co-dependency occurs between the LCR and active gene domains in the YAC transgene.

## Discussion

Intergenic transcription is a widespread phenomenon. Aside from the handful of individual gene loci in which intergenic transcripts have been recognized and studied, it is now apparent from various transcriptome studies that most of the sequence of a mammalian genome is transcribed at one time or another, and that the vast majority of these transcribed regions are non-coding [37]. Many of these non-coding transcripts overlap known coding transcript regions and can be sense or antisense. In many cases, the non-coding transcripts correlate with activity of nearby genes and appear to be regulated in a similar fashion. In other cases, especially where antisense transcripts are involved it has been proposed that these transcripts may play a repressive role. Here we present the characterization of the δβ intergenic promoter, which initiates sense intergenic transcription toward the downstream *HBD* and *HBB* genes. We show that deletion of a 300 bp sequence that contains the δβ promoter results in dramatic alterations to the normal active histone modification profile over a 30 kb downstream region in YAC transgenic mice. As a result, expression of the adult-specific *HBD* and *HBB* genes is severely compromised in erythroid tissues. Our data show that intergenic promoter activity is knocked down by insertion of *loxP* sites flanking the δβ promoter, with a corresponding knock down of adult gene expression. It is not clear why the inserted *loxP* sites cause this knockdown of promoter activity. One possibility is that the upstream *loxP* site disrupts a critical *cis* element needed for full intergenic promoter activity. A search for potential transcription factor binding sites indicates that the upstream *loxP* sequence disrupts a potential E-box binding site, and could potentially bind TAL1 and/or E47. Transgenic line FX14, which has only one *loxP* site in the downstream position, does not suffer from a decrease in intergenic promoter activity, further implicating the upstream site. However, deletion of the region containing this putative binding site results in a doubling of the number of transfected cells that express the EGFP reporter gene in stably transfected K562 cells (compare *Bst*XI and *Apo* in Figure 1B). This is seemingly at odds with the transgenic data but is important to point out that K562 cells express embryonic and fetal globins, HBE and HBG, and do not appear to initiate sense transcripts from the δβ promoter [38]. In combination, these data suggest that this sequence element may be of importance in regulating δβ intergenic promoter activity by repressing transcription in primitive erythroid cells and promoting it in definitive cells.

Our results demonstrate that transcription through the region downstream of the δβ promoter, or the non-coding RNAs produced, or both, play a role in the activation and/or maintenance of the active chromatin structure of the domain. We cannot rule out the possibility that the region also contains an enhancer needed for gene activation, though this seems less likely as it would not account for the wider changes in chromatin structure. The process of transcription itself could potentially affect chromatin structure in a variety of ways. The simplest explanation is that passage of the polymerase results in a transient unfolding of the higher-order chromatin fiber and partial disassembly of nucleosomes. This may create an accessibility window in which, for example, sequence specific binding proteins jump onto their cognate sites [39]. Similarly, disruption may also increase access for chromatin remodelling complexes or histone modification enzymes. Transcription coupled disassembly may also provide the opportunity to insert variant histones such as H3.3 into the transcribed region via the replication independent chromatin assembly pathway [40]. The finding that histone acetyltransferases and methyltransferases are associated with the elongating form of the polymerase, also suggest a possible role for transcription in propagation of modified chromatin domains [41]. Finally, transcription coupled disruption could alter the binding of repressive protein complexes such as the Polycomb group proteins which are thought to be displaced from PREs by intergenic transcription, thus acting as an epigenetic switch [42].

The enormous complexity of the emerging RNA world suggests that the non-coding RNAs themselves also have a role to play in gene control. Some of the most well characterized functional non-coding RNAs, such as *Xist* and *Air* are repressive in nature; however more recent findings have indicated enhancer roles for non-coding RNAs [43]. Non-coding RNAs in the *Drosophila* dosage compensation complex are involved in 2-fold up-regulation of genes on the male X chromosome [44]. The production of non-coding RNAs in the *Hox* clusters have also been tightly correlated with modified chromatin domains [11, 12]. It is possible that epigenetic control of the human β-globin locus involves both transcriptional and non-coding RNA mediated mechanisms since neither is mutually exclusive.

Intergenic transcription could also potentially play a role in nuclear positioning of the beta-globin locus. Studies on the mouse β-globin locus have shown that activation of the locus during erythroid differentiation is LCR independent and involves re-positioning of the locus away from centromeric heterochromatin in correlation with histone hyperacetylation [45, 46]. The fact that histone modification of the human locus is dependent on intergenic transcription, suggests that intergenic transcription may also correlate with nuclear re-positioning. For example, intergenic transcription through the endogenous β-globin locus, which in some cases initiates hundreds of kb upstream [10, 47] could keep the locus tethered to a transcription factory which may facilitate near-continuous, high-level transcription of the active globin genes [48, 49], while at the same time maintaining a heritable epigenetic memory of the active state. A role for the LCR in contributing to promotion or maintenance an active chromatin profile in the locus may also be proposed. Dostie et al., [50] demonstrated a long-range interaction with the δβ promoter region in human erythroid cell line, suggesting that the LCR may stimulate intergenic transcription that, in turn, may promote an active state. Other sequences in the γδ region between the ^A^γ-globin gene (HBG2) and the δ-globin gene (HBD), including the *HBBP1* pseudogene (ψβ-globin) and the *BGLT3* non-coding gene that is between HBG2 and HBBP1, contribute to long range interaction [51, 52] Our data show that the δβ promoter region is a novel *cis*-regulatory element that is well downstream of HBBP1 pseudogene, but upstream of the HBD gene.

A note of caution must be added to these studies since the locus under study is a transgene. Incongruent results have often been obtained when comparisons are made between mutations in the human β-globin transgene locus and seemingly similar mutations in the endogenous mouse β-globin locus. Part of this may be due to the well-documented fact that human transgenes in mouse erythroid cells do not exactly recapitulate the regulatory pattern of the endogenous locus. Our analysis of *ex-vivo* erythroid precursors from patients with a naturally occurring deletion of the δβ promoter helps to illustrate this point [19]. The Corfu thalassemia mutation is a 7.2 kb deletion that removes the δβ promoter and the 5’ end of the *HBD* gene. Unlike the situation in transgenic mice, the deletion does not lead to a general shut down of the affected globin locus and LCR. Instead we found that intergenic transcripts appeared to initiate from an intergenic promoter upstream of the fetal-specific *HBG* genes. This correlated with an open chromatin structure over the *HBG* genes and the downstream *HBB* gene, and high-level transcription of both *HBG* genes and *HBB* in the adult cell environment. It is tempting to speculate that binding sites for BCL11A or the PYR repressor complex, factors implicated in *HBG* silencing, are deleted in the larger Corfu mutation [53, 54], which may partly explain the differences in *HBG* silencing.

## Material and Methods

### Human primary erythroid cell culture

Samples of venous blood (50 ml) were collected from informed and consenting volunteers using EDTA vacuette tubes and spun on Histopaque-1077 (Sigma) to isolate the mononuclear cells. Following six PBS platelet removal washes these were resuspended at 1×10^6^ cells/ml in phase one media: Stemspan medium (Stemcell Tech), 1μg /ml cyclosporin A (Sigma), 20 ng/ml IL-3 (Peptrotech), 20 ng/ml IL-6 (Peptrotech) and 50 ng/ml stem cell factor (Miltenyi Biotech). These were incubated for 24 hours at 37°C; non-adherent cells were transferred to a new flask and incubated for a further 6 days at 37°C in 5% CO_2_. The cells were washed in PBS and then seeded at 1×10^5^ cells/ml in phase two media: Stemspan medium, 2U/ml EPO (Espex), 5 ng/ml IL-3, 2×10^−6^ M dexamethasone (Sigma), 10^−6^ M β-estradiol (Sigma) and 20 ng/ml stem cell factor. Cells were collected at different time-points of phase two growth to monitor erythroid differentiation and extract RNA using Tri-reagent (Sigma). Differentiation was assessed using qRT-PCR (with Taqman probes for HBB and HBG2, and GAPDH to normalise) and May-Grunwald morphological staining (data not shown).

### Analysis of RNA extracted from phase two erythroid culture

Microarrays were designed by Nimblegen Roche to cover the β-globin locus at high resolution. 100 ng of RNA was treated with DNase (Sigma) before being converted to cDNA using the Transplex complete whole transcriptome amplification kit (Sigma) as per the manufacturer’s guidelines. The cDNA was RNase (Sigma) treated and cDNA precipitated and labelled using the Nimblegen Expression Array kit using the manufacturer’s protocol. The labelled cDNA was hybridised to a single chamber in a Nimblegen 12plex microarray slide following the manufacturer’s guidelines. The slide was washed and scanned at 3 mm resolution using an InnoScan 700 Microarray Scanner and TIFF images were generated using Mapix ver 5.1 software. The TIFF images were aligned and converted into probe intensity values using the Nimblegen DEVA software with background correction and uploaded onto UCSC genome browser.

To confirm the peak of transcription over the δβ-promoter region in human erythroid cells quantitative RT-PCR was performed; cDNA was first generated from 150 ng of DNase-treated (Sigma) RNA using the High Capacity RNA-to-cDNA kit (Applied Biosystems). Quantitative PCR was carried out on the BioRad CFX96 QPCR machine using IQ SYBR green supermix (Bio-Rad), 0.5 μM concentration of each primer and 0.9 μl of cDNA. No-RT controls and no-template controls verified the lack of genomic or PCR contamination respectively (not shown). Standard curves for the U-amplicon and D-amplicon were generated by amplifying genomic DNA at a range of concentrations and used to quantify the relative levels of transcription in the region. The ratio of downstream over upstream signal was used to estimate the changing transcriptional pattern during erythroid differentiation.

Primer sequences:

Downstream-F (forward primer): 5’-ATGCCAATGTGGGTTAGAATG-3’

Downstream–R (reverse primer): 5’-CATACCATGTGGCTCATCCTC-3’

Upstream-F (forward primer): 5’-TTAAATGCCAGTGCTCTCCAC-3’

Upstream-R (reverse primer): 5’-CAGAAGGAAACTAGGATGTGTCC-3’

For strand-specific RT-PCR of the D and U amplicons cDNA was generated from 150 ng of DNase-treated RNA using the GoScript™ Reverse Transcription System (Promega) with either the Downstream-F or Downstream-R primer. PCR was then performed using Downstream-F and Downstream-R primers. To assess the RNA integrity an additional cDNA reaction was performed by using random-primers in the RT step.

### YAC transgenic mice

264W transgenic mice carrying a single copy of a 150 kb human β-globin locus YAC were previously described [32]. A 213 kb YAC containing the human β-globin locus [28] was used to generate the rest of the transgenic lines. The YAC was modified using the pop-in-pop-out recombination procedure [55] to introduce *loxP* sites upstream and downstream of the minimal δβ intergenic promoter. Briefly, a gene targeting vector was generated by inserting *loxP* sites at the *Bse*RI and *Cla*I restriction sites of the 2.5 kb *Bgl*II restriction fragment (see Figure 3). The modified YACs were then used to generate transgenic mice as previously described [26]. The structural integrity of the β-globin loci in the obtained transgenic animals was analysed as described in [27]. Briefly, *Sfi*I-digested DNA was fractionated by pulsed-field gel electrophoresis and Southern blots were prepared. The membrane was cut into strips comprising individual gel lanes. Each strip was hybridized with a single probe. Probes utilised include: probe 1, 0.7 kb *Pst*I fragment upstream of HS3; probe 2, 1.9 kb *Hind*III fragment upstream of HS2; probe 3, 3.7 kb *Eco*RI fragment spanning the *HBE* gene; probe 4, 2.4 *Eco*RI fragment downstream of the *HBG1* gene; probe 5, 1.9 kb *Xba*I fragment downstream of the beta-like pseudogene; probe 6, 2.1 kb *Pst*I fragment upstream of the *HBD* gene; probe 7, 0.9 *Eco*RI-*Bam*HI *HBB* fragment; probe 8, 1.4 kb *Xba*I DF10 3’HS1 fragment; probe 9, 1.9 kb *Bgl*II HPFH-3 fragment; probe 10, 0.5-kb *Hind*III H500 fragment; probe 11, 1.5-kb *Eco*RI-*Bgl*II HPFH-6 fragment (all described in [27]). Transgene copy numbers were estimated by quantitative PhosphorImager analysis of Southern blots. To generate a line with a deletion of the δβ intergenic promoter, mice were bred to Deleter Cre recombinase-expressing mice [31]. After several generations, animals carrying a single, complete copy of the human β-globin locus with a deletion of the δβ promoter were identified. The integrity of the transgene after recombination was confirmed by pulsed-field gel electrophoresis of *Sfi*I-digested DNA and subsequent Southern blotting and hybridisation with probe D, 2.5 kb *Bgl*II fragment spanning the δβ promoter. Deletion of the δβ promoter and copy-number reduction were verified by Southern hybridization of *Bgl*II-digested genomic DNA with probes D and *Thy1*, followed by PhosphorImager quantification.

### Reporter constructs, transfection of cultured cells, and FACS analysis

50 ng of mouse genomic DNA from lines 264W, FX115 and Δ115 were amplified by PCR with primers 5’-CTTGAGTGCATTGACAAAATTACC-3’ and 5’-CAGACTTGGACCATGACGGTG-3’. The PCR products were cloned into pGEM-T easy (Promega) and the sequence of the resulting plasmids was verified. The δβ promoter fragments were released by *Eco*RI/*Cla*I (264W) or *Eco*RI/*Sal*I (FX115 and Δ115) digestions, blunted with T4 DNA polymerase (New England Biolabs) and ligated into vector pEGFP-C1 (promoterless EGFP, Clontech) at the *Sma*I site. Plasmids were verified by sequencing.

Cells were grown in RPMI 1640 supplemented with 10% foetal calf serum and Penicillin/Streptomycin. 5×10^7^ cells were transfected by electroporation with 50 μg of linearized plasmid DNA at 350 V, 1000 μF in 500μl of 1x Hepes-Buffered Saline. Plasmid pEGFP-N1 (Clontech), expressing EGFP under the control of a CMV promoter, was also used as a positive control. After transfection cells were transferred to non-selective medium for 24 hours followed by 14 days in medium with 600 μg/ml G418. EGFP expression was analysed by FACS performed on a FACSCalibur. Background was determined by analysing non-transfected cells and percentage of green cells as well as mean fluorescence intensity was determined using the CellQuest software.

### RNA FISH

RNA FISH was performed as previously described [9]. Cocktails of hapten-labelled oligonucleotides (Oswel) were used as probes, sequences provided below:

Dinitrophenol-labelled human *HBE* intron 2 probes:

5’-AATCTTGAGGACTTTCCCAATCAACTTGCT-3’

5’-GTTCTAACATCAAGCTCGACCCATGATTTC-3’

5’-CCTTCACTTGGTCACAATACTGATGGGAAA-3’

5’-CTGACCTCAAACTGTTCCAAGGTTTGTGCC-3’

Digoxigenin-labelled human *HBG* intron 2 probes:

5’-CGACCTGGACTTTTGCCAGGCACAGGGTCC-3’

5’-TCACTCCCAACCCCAGTATCTTCAAACAGC-3’

5’-GCATCTTTTTAACGACCATACTTGTCCTGC-3’

5’-ACAGAGCTGACTTTCAAAATCTACTCCAGC-3’

Dinitrophenol-labelled human *HBB* intron 2 probes:

5’-TTCCACACTGATGCAATCATTCGTCTGTTT-3’

5’-TGTGTACACATATTAAAACATTACACTTTA-3’

5’-ATTAGCAATATGAAACCTCTTACATCAGTT-3’

5’-AGTAATGTACTAGGCAGACTGTGTAAAGTT-3’

Dinitrophenol-labelled mouse *Hbb-b1* intron 2 probes:

5’-TCAACACTCCACACACAGTCATGGAGACTG-3’

5’-GATATCAGGATGGGAAGTAAATAACCAGCT-3’

5’-AAACAAACGTAGATCAAAGAAGAAGAAATG-3’

5’-AGCTATGAGAAGAAACAGGGACATATCTTC-3’

Digoxigenin-labelled human *HBB* exon 3 probes:

5’-TTTAATAGAAATTGGACAGCAAGAAAGCG-3’

5’-AGGCCCTTCATAATATCCCCCAGTTTAGTA-3’

Dinitrophenol-labelled probes were detected with Texas Red-labelled antibodies (primary antibody from Serotec, secondary antibodies from Jackson ImmunoResearch); Digoxigenin-labelled probes were detected with FITC-labelled antibodies (primary antibody from Roche, secondary antibodies from Jackson ImmunoResearch). Slides were examined on an Olympus BX41 epifluorescence microscope. To determine the percentage of cells with human transgene signals, a minimum of 300 loci were counted for each data point.

### RT-PCR

Total RNA was isolated from adult anaemic spleen 5 days after phenylhydrazine injections [56] using the RNeasy mini kit (Qiagen) according to the manufacturer’s instructions. RNA samples were DNase I-treated with 1U of RQ1 DNase (Promega) per μg of RNA and purified through a second RNeasy column. 1 to 5μg of total RNA was reverse-transcribed with Superscript III reverse transcriptase (Invitrogen) following manufacturer’s protocol at 50°C in the presence of 2 U/μl RNasin (Promega). RT negative controls in which the reverse transcriptase enzyme was omitted were set up in parallel. Quantitative RT-PCR was performed by real-time PCR in an ABI PRISM 7000 Sequence Detection System using SYBR green PCR Master Mix (Applied Biosystems). All PCR reactions were carried out in duplicates and carbonic anhydrase II was used as a normalisation control. Serial dilutions of genomic DNA (primary and intergenic transcripts) or of a plasmid containing the carbonic anhydrase II cDNA PCR product were used as standard curves for absolute quantification.

The following primers were used for PCR amplification of cDNA:

*HBD* gene, intron 2 forward primer: 5’-TGGGGATCAGTTTTGTCTAAGATTTGGG-3’

*HBD* gene, intron 2 reverse primer: 5’-GCGGAGAAGAGGTAGGCAGATACATGC-3’

*HBB* gene, intron 2 forward primer: 5’-GACGAATGATTGCATCAGTGTGGAAGTC-3’

*HBB* gene, intron 2 reverse primer: 5’-TGCGGAGAAGAAAAAAAAAGAAAGCAAG-3’

intergenic RT3B forward primer: 5’-TCATCTGGGATTTTGGGGACTATGTC-3’

intergenic RT3B reverse primer: 5’-CAGACTTGGGAATTGGGATTATACAGGC-3’

intergenic RT4 forward primer: 5’-CTTGTGTTCACGACTGACATCACCG-3’

intergenic RT4 reverse primer: 5’-GGAGACAAATAGCTGGGCTTCTGTTG-3’

mouse carbonic anhydrase II forward primer: 5’-GAGTTTGATGACTCTCAGGACAATGCAG-3’

mouse carbonic anhydrase II reverse primer: 5’-AGATGAGCCCCAGTGAAAGTGAAAC-3’

### Chromatin immunoprecipitation

Histone modification profiles across the human β-globin locus were assessed by native chromatin immunoprecipitation [57] and performed exactly as described by [10]. Briefly, single-cell suspensions of erythroid cells were resuspended to 2 x10^7^ cells/ml in ice cold RSB buffer (10 mM Tris-HCl, pH 7.5, 10 mM NaCl, 3 mM MgCl_2_), 0.1% Triton X-100, 0.5 mM DTT, 0.1 M sucrose, 0.1 mM PMSF, 5 mM Na butyrate, supplemented with protease inhibitor cocktail (Sigma). The cells were homogenised in a cold B-type Dounce homogeniser and diluted with an equal volume of the same buffer with 0.25 M sucrose. The suspension was layered onto a sucrose cushion of 0.33 M sucrose, 5 mM MgCl_2_, 10 mM Tris pH 8.0, 0.5 mM DTT, 0.1 mM PMSF, 5 mM Na butyrate. This was then centrifuged at 800 × g for 5 min at 4°C to pellet nuclei. Chromatin was digested with micrococcal nuclease generating DNA predominantly mononucleosomal in length. ChIP was carried out using the following rabbit polyclonal antibodies: anti-trimethyl-histone H3 (K4) (Abcam), anti-dimethyl-histone H3 (K4) (Upstate Biotechnology), anti-acetyl-histone H3 (K9/K14) (Upstate Biotechnology).

The representation of multiple sequences across the human β-globin locus in the Input and Bound fractions was determined by quantitative real-time PCR. Real-time PCR was performed in an ABI PRISM 7000 Sequence Detection System using SYBR green PCR Master Mix (Applied Biosystems). All PCR reactions were carried out in duplicates using 3 ng Input or Bound DNA as template. For primer sequences, see Miles et al. [10]. The ratio of Bound to Input DNA was calculated using the comparative C_T_ method, Bound/Input = 2^(Input Ct – Bound Ct)^ [58]. Data were normalised to the most 3’ data point, downstream of 3’HS1, which shows low enrichment for all samples.

## Acknowledgments

We thank L. Mercer, M. Anderton, A. Nanou, and D. Dimitrova for their assistance. This work was supported in part by the Medical Research Council and the Biotechnology and Biological Sciences Research Council, UK, by USPHS NIH grants DK053510, HL067336, and HL111264 awarded to KRP and by a grant from Sparks awarded to DRFC.

